# Brain Metabolic Network Covariance and Aging in a Mouse Model of Alzheimer’s Disease

**DOI:** 10.1101/2023.06.21.545918

**Authors:** EJ Chumin, CP Burton, R Silvola, EW Miner, SC Persohn, M Veronese, PR Territo

## Abstract

**INTRODUCTION:** Alzheimer’s disease (AD), the leading cause of dementia worldwide, represents a human and financial impact for which few effective drugs exist to treat the disease. Advances in molecular imaging have enabled assessment of cerebral glycolytic metabolism, and network modeling of brain region have linked to alterations in metabolic activity to AD stage.

**METHODS:** We performed ^18^F-FDG Positron Emission Tomography (PET) imaging in 4-, 6-, and 12-month-old 5XFAD and littermate controls (WT) of both sexes and analyzed region data via brain metabolic covariance analysis.

**RESULTS:** 5XFAD model mice showed age related changes glucose uptake relative to WT mice. Analysis of community structure of covariance networks was different across age and sex, with a disruption of metabolic coupling in the 5XFAD model.

**DISCUSSION:** The current study replicates clinical AD findings and indicates that metabolic network covariance modeling provides a translational tool to assess disease progression in AD models.

**RESEARCH IN CONTEXT:** *SYSTEMATIC REVIEW:* The authors extensively reviewed literature (e.g., PubMed), meeting abstracts, and presentations on approaches to evaluate brain network analysis in animal models. Based on the available data, there were clear gaps in our understanding of how metabolic networks change with disease progression at the preclinical phase, thus limiting the utility of these measures for clinical comparison in Alzheimer’s disease (AD).

*INTERPRETATION:* Our findings indicate that employing metabolic covariance modeling in mouse models of AD and littermate controls of both sexes with age provides a mechanism to evaluate brain changes in network function which align closely with previous clinical stages of AD. Moreover, utilizing open-source clinical tools from the Brain Connectivity Toolbox (BCT), we demonstrated that brain networks reorganize with AD progression at multiple levels, and these changes are consistent with previous reports in human AD studies.

*FUTURE DIRECTIONS:* The open-source framework developed in the current work provides valuable tools for brain metabolic covariance modeling. Such tools can be used in both preclinical and clinical settings and they enable more direct translation of preclinical imaging studies to those in the clinic. When matched with an appropriate animal model, genetics, and/or treatments, this study will enable assessment of *in vivo* target engagement, translational pharmacodynamics, and insight into potential treatments of AD.

## INTRODUCTION

Alzheimer’s disease (AD) represents the leading cause of dementia worldwide, and currently there are few approved therapies to slow, prevent or reverse the disease process [1, 2]. The number of people affected by AD in the US is expected to reach 14 million by 2050, which translates into a financial burden that will exceed 9 trillion dollars worldwide [3]. The molecular characteristics of AD are extracellular β-amyloid (Aβ) plaque formation, and accumulation of hyperphosphorylated tau in neurofibrillary tangles. These characteristics lead to inflammation, synaptic impairment, and cognitive decline [1, 2]. Anatomical investigations with magnetic resonance imaging (MRI) have revealed atrophy as a consequence of AD neuropathology [4]. Additionally, molecular imaging with positron emission tomography (PET) of amyloid, tau, and glycolytic metabolism [5], have demonstrated a clinical utility for neuroimaging in predictive/clinical diagnosis and care for AD and related dementias. Several prior studies, in both nonhuman and human samples, have shown altered regional metabolic demands in AD, as measured with ^18^F-2-fluorodeoxyglucose (^18^F-FDG) PET [6-9], where AD was associated with regional hypometabolism in the brain. By contrast, in early probable AD stages (referred to as asymptomatic preclinical or at-risk stages), regional hypermetabolism has been shown [10, 11]. Furthermore, recent work suggests that AD progression clinically is linked to alterations in metabolic activity of brain networks (default mode network and AD-related pattern) measured by ^18^F-FDG PET [12].

In addition to measures of regional alterations with AD, inter-regional relationships have been investigated though applications of graph theory and network neuroscience (for review, see Bassett and Sporns [13]). In the context of ^18^F-FDG PET, these relationships are often referred to as metabolic connectivity or metabolic covariance networks [14] and can be quantified as inter-subject covariance networks (cross-correlation of regional values across a homogenous sample or group) [15]. Such networks have been observed in healthy older adults, and are in-part related to functional connectivity networks [16]. Alterations in metabolic covariance have also been reported in multiple degenerative diseases [15, 17-20]. Among them, the AD group was shown to have altered metabolic connectivity [14], including increased entropy, connection strength, and clustering coefficient [15].

In preclinical rodent models of AD, there is a lack of consensus about characteristic glycolytic metabolism among ^18^F-FDG PET studies (for review see [21]). The 5XFAD mouse model, which expresses five mutations of human familial AD genes [22], has been shown to have both increased [23] and decreased ^18^F-FDG-measured glucose metabolism [8, 24]. Variability in outcomes is likely due to methodological differences, notably whether the animals were anesthetized, fasted, and how tracer uptake was handled. Regional analysis has typically utilized standardized uptake values (SUV), which assume that the total volume of distribution of the tracer is related to the body mass and seek to normalize the regional distribution by this factor. Alternatively, SUV ratios (SUVR) have also been used, which represent the uptake as a ratio of the region of interest to a reference region (i.e., cerebellum or corpus callosum), thus minimizing dose and noise effects. In addition, outcome variability may also be due to underlying genetics and pathological fluctuations in cerebral metabolism in AD across measurements. Collectively, these studies reinforce the importance of *in-vivo* imaging of AD neuropathology and suggest that analysis of glycolytic metabolism via interregional covariance may provide valuable insights into AD progression. In this study, we explored whether a metabolic covariance structure in ^18^F-FDG-derived networks from mice differed in 5XFAD compared to wild-type (WT) animals with respect to sex and age. We hypothesized that 5XFAD mice cohorts will display characteristics indicative of neurodegeneration and disrupted glucose update/utilization in their networks compared to their WT counterparts.

## METHODS AND MATERIALS

### 5XFAD Transgenic Mouse Model

5XFAD mice overexpress five familial Alzheimer’s Disease (FAD) mutations: the APP (695) transgene includes the Sweden (K670N, M671L), Florida (I716V), and London (V717I) mutations, and the PSEN1 transgene accounts for the M146L and L286V FAD mutations. The 5XFAD line was made congenic on the C57BL/6J background in 2011 at Jackson Laboratories to minimize concerns related to allele segregation and the high variability of the original hybrid background. 5XFAD mice exhibit cerebral amyloid deposition, gliosis, and progressive neurodegeneration accompanied by cognitive and motor deficiencies, recapitulating many of the features of human AD. However, studies have shown that neurofibrillary tangles (NFTs) are not typically present in the 5XFAD model [25], indicating a key distinction between AD pathology in human disease and this model.

### Animal Housing

Male hemizygous 5XFAD mice (MMRRC stock #: 34848) were crossed with female C57BL6/J mice (JAX stock #: 000664) to maintain the 5XFAD and non-transgenic WT colonies at Indiana University. Up to five mice were housed per cage with Aspen SaniChip bedding and ad libitum LabDietR 5K52/5K67 (6% fat) feed. The colony room was kept on a 12:12 schedule with the lights on from 7:00 AM to 7:00 PM daily. Transgenic and WT mice were initially ear-punched for identification; following genotyping, mice were identified with p-chip (PharmaSeq) microchips placed at the base of the tail. Both male and female littermate mice aged 4, 6, and 12 months were used for this study. These ages reflect early disease state (4 months), moderate disease state (6 months), and late disease state (12 months). In all cases, non-transgenic littermates were used as WT controls. All studies were approved by the Institutional Animal Use and Care Committees (IACUC) at Indiana University School of Medicine.

### Magnetic Resonance Imaging (MRI)

High-contrast gray matter images were acquired at least 2 days prior to PET imaging. Mice were anesthetized (induced with 5% isoflurane in medical oxygen, maintained with 1–3% isoflurane) prior to MR image acquisition. High-resolution T2-weighted (T2W) MRI images were acquired using a 3T Siemens Prisma MRI scanner (Singo, v7.0), outfitted with a 4-channel head coil, anesthesia, and bed system (RAPID MR, Columbus, OH, United States). SPACE3D images were acquired with the following parameters: TA: 5.5 min; TR: 2080 ms; TE: 162 ms; ETL: 57; FS: On; Average: 2; Excitation Flip Angle: 150; Norm Filter: On; Restore Magnetization: On; Slice Thickness 0.2 mm; Matrix: 171 x 192; and FOV: 35 x 35 mm, yielding 0.18 x 18 x 0.2 mm resolution images.

### ^18^F-FDG Positron Emission Tomography (PET) Image Acquisition

Prior to tracer administration, mice were fasted for a minimum of 12 hours to effectively normalize serum glucose levels for all mice in an imaging cohort. Mice were injected intraperitoneally (IP) with 3.7 to 9.25 MBq (100 to 250 μCi) of ^18^F-FDG, with the final volume not exceeding 10% of the animal’s weight. After injection, mice were placed in their home cage for 45 minutes to allow for tracer uptake [22]. The mice were then anesthetized with 5% isoflurane gas, positioned on the imaging bed, and scanned on the IndyPET3 scanner [26]. During acquisition, anesthetic plane was maintained with 1-3% isoflurane. Post listmode data collection, PET images were calibrated, decay and scatter-corrected, and reconstructed into a single-static image volume with a 60 mm field of view using filtered-back-projection [27], with an effective resolution of 1.1 x 1.1 x 1.2 mm (1.45 mm^3^). Images were converted from NetCDF to an Analyze 12-compatible format prior to analysis.

### Image Preprocessing

PET and MRI images were co-registered to stereotactic mouse brain coordinates [28] using a rigid-body mutual information-based normalized entropy algorithm with 9 degrees of freedom implemented in Analyze 12 (AnalzeDirect, Stilwell, KS, United States). After registration, 56 bilateral regions were extracted. Standard uptake value ratios (SUVRs) to a constant cerebellar reference region to control for dose and cerebral metabolic confounds were computed as:

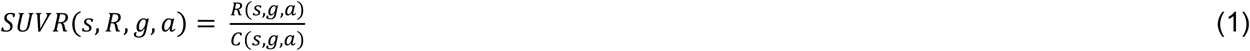

where the SUVR for a given animal (*s*), region (*R*), genotype (*g*), and age (*a*) is the ratio of the region of interest activity *R(s,g,a)* and cerebellar reference region activity *C(s,g,a)* in Bq/ml. To minimize the likelihood of partial volume effects, region volumes were selected where the smallest region (i.e., fornix = 178 mm^3^) exceeded the effective scanner sample volume by ≥123-fold. At the limit, using a partial volume factor of 5 voxels, the smallest resolvable effective volume would be 7.25 mm^3^, and is well within the smallest region volume used in the current study. After SUVRs were calculated and registered, left and right brain regions were averaged on the assumption of bilateral symmetry.

### Covariance Network Analysis

Network analysis was computed in Matlab (MathWorks, version 2021b; Natick, MA) with the use of the Brain Connectivity Toolbox (BCT) [29], generalized Louvain modularity [30], and hierarchical consensus clustering [31]. Analysis code is publicly available in the CovNet package (https://github.com/echumin/CovNet).

### Network Estimation and Thresholding

Covariance networks were computed for 3 age groups (4, 6, and 12 months) of WT (males: n=12, 11, 9; females n=12, 11, 10) and 5XFAD mice (males: n=11, 10, 9; females: n=12, 12, 10), using Pearson correlation of regional *z*-scored SUVR values across animals within group. The resultant covariance network for each group is then a region-by-region (27 x 27 region) adjacency matrix that indicates degree of similarity of ^18^F-FDG quantified metabolic coupling among all region pairs. Three network variants were generated: 1) weighted unthresholded; 2) weighted thresholded at *p* < 0.05 correlation significance; and 3) weighed thresholded at *p* < 0.01. Edge-level correlation significance was assessed with a random shift null model, where SUVR values were permuted independently for each region (i.e., shuffling animals within group), destroying the covariance structure with 10,000 permutations. The distribution of 10,000 null covariance networks was then used to estimate permutation p-values at each edge as the fraction of times the empirical correlation exceeded the null in magnitude (positive and negative correlations were handled separately). Modularity maximization was performed on unthresholded networks as they contain the most information, while both unthresholded and thresholded networks were used to assess network properties. Thresholds below *p* < 0.01 were not investigated as they produced fractured networks (where a node or a group of nodes are disconnected from the rest of the network) in more than half of the groups.

### Global and Nodal Network Measures

The following network measures were computed on thresholded networks using the BCT: network density – number of connections, node degree – number of connections for each region, positive and negative nodal strengths – sum of the positive or negative covariance values for each region, clustering coefficient – number of fully connected triangles around a node taking into account the weight and sign of the connections (coefficient type 3 in the BCT clustering_coef_wu_sign function) [32]. Two variants of hub regions were also identified as the top quartile (7 regions) in network strength (strength-based) (cumulative strength computed as the sum of the absolute values) or degree (degree-based). Strength-based hubs were estimated from unthresholded and thresholded networks, while degree-based hubs were estimated from thresholded networks only.

### Multiresolution Consensus Clustering

Community detection, also referred to as modularity maximization, is the process of identifying a partition of nodes in a network into groups or communities, such that within community connectivity is greater than between community. However, this raises an issue of scale, asking what is the optimal number of communities in a network? While there is no correct answer to this question, the Multiresolution Consensus Clustering (MRCC) [31] approach addresses this by identifying communities across spatial scales, and then quantitatively identifying a consensus partition from the set. Here MRCC was applied to weighted unthresholded networks, generating a set of 10,000 initial partitions. From this set of partitions, MRCC then computes a co-assignment matrix (i.e., probability that any node pair is in the same community), a dendrogram of partitions across spatial scale, and a consensus partition given an alpha threshold (α = 0.05). The derived consensus partitions were then qualitatively compared and used to investigate group differences in mean module/community SUVR values.

### Statistical Analysis

Edge weight distributions of networks as well as distributions of nodal measures (i.e., degree, strength, and clustering coefficient) were compared between groups via 2-sample Kolmogorov Smirnov (KS) tests. Pairwise edge-level differences in group covariance networks were assessed with permutation *t*-tests (10,000 group label permutations on regional SUVR values, recomputing a null covariance network at each permutation). Multiple comparisons correction for number of edges (i.e., connections) was done with a false discovery rate (FDR) adjusted at *q*<0.05. Module average SUVR values were extracted from all groups using a single groups partition as reference, so that the same regions are being averaged for all groups. An n-way analysis of variance (Matlab) with, age, sex, and genotype as factors, was used to test for main effects and two-way interactions. In the presence of significant effects, post hoc multiple comparisons were assessed with Tukey-Kramer.

## RESULTS

### Characterization of Group Covariance

Metabolic covariance matrices and distributions of edge weights (i.e., Pearson correlations) are shown in Figure 1. WT covariance matrices were qualitatively more structured compared to 5XFAD, notably in males. Pairwise KS tests comparing male versus female networks within genotype showed that only the WT 6-month network-derived edge weight distributions were significantly different (Figure 1B, KS statistic 0.14, *p* < 0.005). Thresholded covariance networks for *p’s* < 0.05 and 0.01 are shown in Figures S1 and S2, respectively. Between WT and 5XFAD, distributions of edges weights were not significantly different, with the exception of females at 12 months (Figure 2F, KS statistic 0.1, *p* < 0.05). Edgewise group comparisons were carried out with permutation testing, with edges below *p* < 0.05 shown in Figure 2. No individual edges survived FDR correction. In male mice at 4 months, at *p* < 0.05 uncorrected significance, 4 regions stand out as having altered coupling to a large portion of the other regions in the network: the ventral, medial, and lateral orbitofrontal cortices, and the fornix (Figure 2A). Differences in females at 4 months identified the agranular insular and dysgranular insular cortices with the most differences to the rest of the network (Figure 2D). At 6 months of age for both sexes, some edgewise differences were identified, but no regions showed differences to a notable portion of the rest of the network (Figure 2B, E). Finally, at 12 months, multiple regions showed differences with ∼20% of the network (i.e., secondary somatosensory, retrosplenial dysgranular, perirhinal and perilimbic cortices, hippocampus, fornix, auditory and frontal association cortices; Figure 2C), while in females only the ventral orbital cortex stood out (Figure 2F).

**Figure 1.**
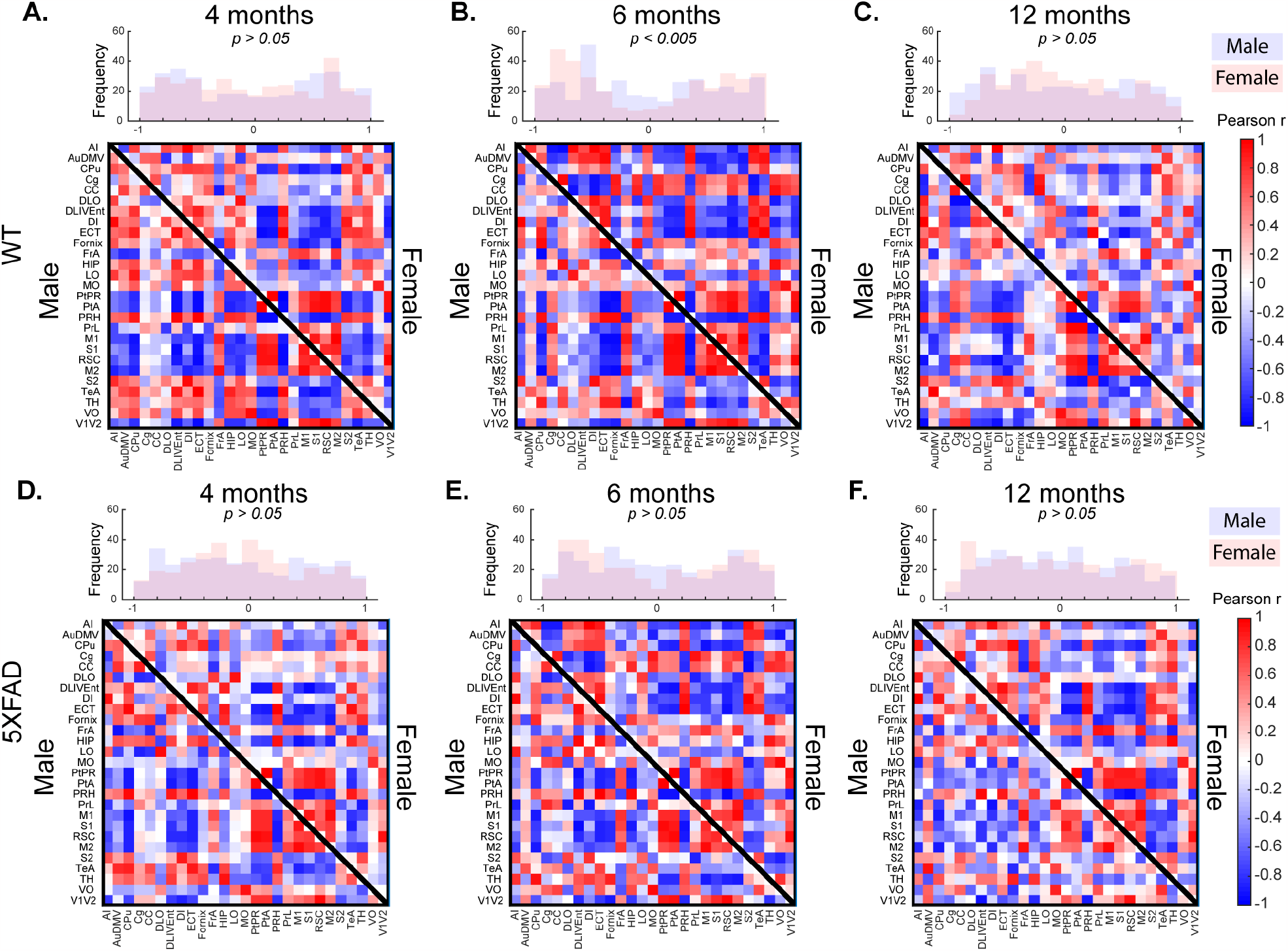
Metabolic (^18^F-FDG PET) covariance matrices for (A-C) wild-type (WT) and (D-F) 5XFAD groups. Histograms show the correlation coefficient distributions from male (lower triangle of matrix) and female (upper triangle of matrix) covariance matrices at 4 months (A, D), 6 months (B, E), and 12 months (C, F) old. Covariance was computed as Pearson correlation of *z*-scored regional SUVR values across animals within each group. Full names for row/column region labels can be found in Table S1. For visualizations of thresholded networks see Figure S1 and S2 for *p* < 0.05 and *p* < 0.01 correlation significance, respectively.

**Figure 2.**
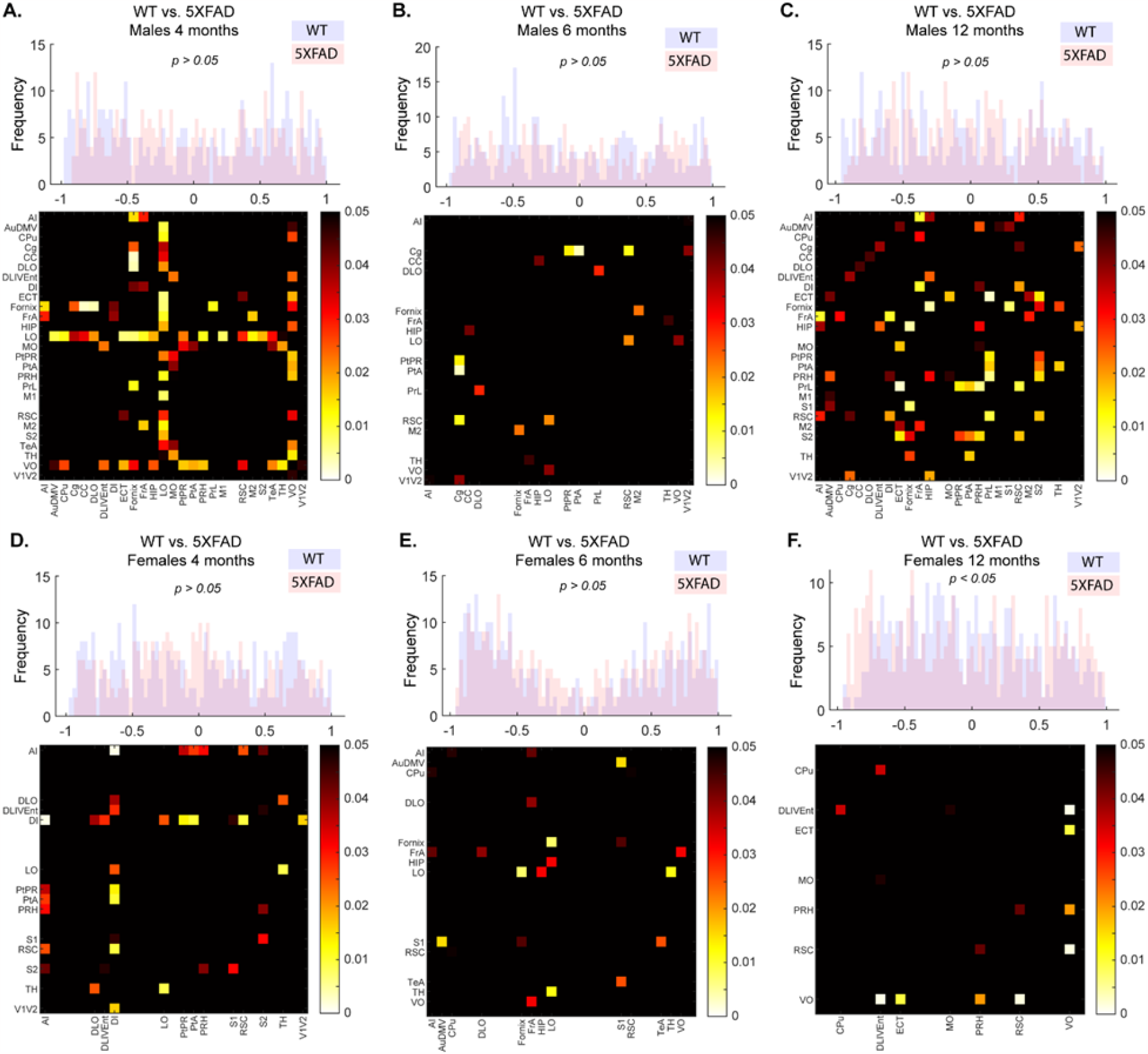
Covariance network comparisons across genotype. Histograms are distributions of network edge weights statistically compared with a Kolmogorov-Smirnov test at *p* < 0.05 threshold. Only in Females at 12 months (F) were the distributions significantly different. Individual edges were tested for group differences with permutation testing (10,000 permutations), with matrices showing uncorrected significance at *p* < 0.05. Multiple comparisons were assessed with FDR, with no adjusted significant edges identified.

Examining the global network properties, network densities (i.e., number of connections) were inversely related to age at both p-value thresholds in WT male mice. Compared to WT, 5XFAD male mice showed a lower network density at 4 months, were similar at 6 months, and were drastically lower at 12 months (Figure 3A-B). Female mice showed an inverted u-shape pattern across age that was similar in both WT and 5XFAD groups, with one distinction; the 5XFAD female mice network had a higher density relative to WT at 12 months (Figure 3A-B). Nodal network properties followed pattern similar to network density, which was expected given the strong impact of network density on other network metrics [29]. Pairwise comparisons across genotype identified differences in distributions of nodal degree (i.e., number of connections; Figure 3C) and positive and negative nodal strength (i.e., sum of positive and negative edge weights, or correlation coefficients, see Figure 3D-E, respectively) in both sexes and multiple age groups.

**Figure 3.**
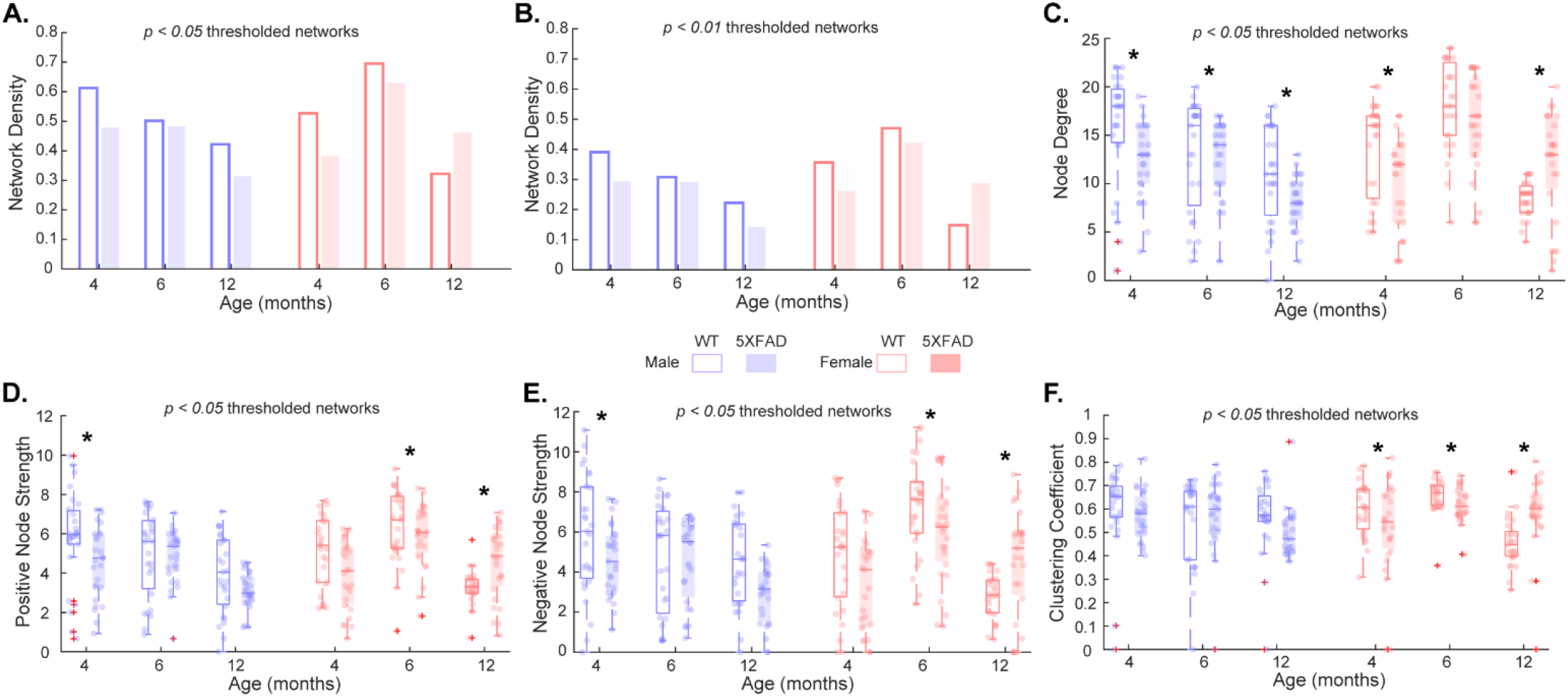
Properties of thresholded covariance networks for wild-type (WT) and 5XFAD groups at 4, 6, and 12 months old. **(A-B)** Network densities (*p* < 0.05 and 0.01). **(C)** Degree, **(D)** positive and **(E)** negative strength, and **(F)** clustering coefficient nodal distributions of *p* < 0.05 thresholded covariance networks. Degree, strength, and clustering coefficient for *p* < 0.01 thresholded networks were lower in magnitude (as expected) but overall similar (Figure S3). *Denotes *p* < 0.05, 2-sample Kolmogorov-Smirnov test significance. +Denote data points that fall outside the 1.5*interquartile range below the 25^th^ and above the 75^th^ percentile.

Differences in clustering coefficient (i.e., measure of network segregation, or how many of the nodes’ connected neighbors are connected to each other) between WT and 5XFAD were only seen in female groups (Figure 3F). At a more restrictive network threshold of *p* < 0.01, a similar pattern was observed (Figure S3).

### Hubs of Metabolic Covariance

While there was some overlap in hub regions overall, uniquely identified hubs showed a divergence by genotype. When hubs were defined based on degree (number of connections, Figure 4), the difference between WT and 5XFAD networks tended to increase proportionate to age. With respect to sex, males tended to express a greater magnitude of alterations in degree hubs compared to females (less overlap/consensus). Females at 6 months, as with above discussed metrics, were similar between WT and 5XFAD, which was not the case for either 4-or 12-month female groups. Alternatively, defining hubs based on strength (sum of weights) implicated a similar set of regions (Figure S4).

**Figure 4.**
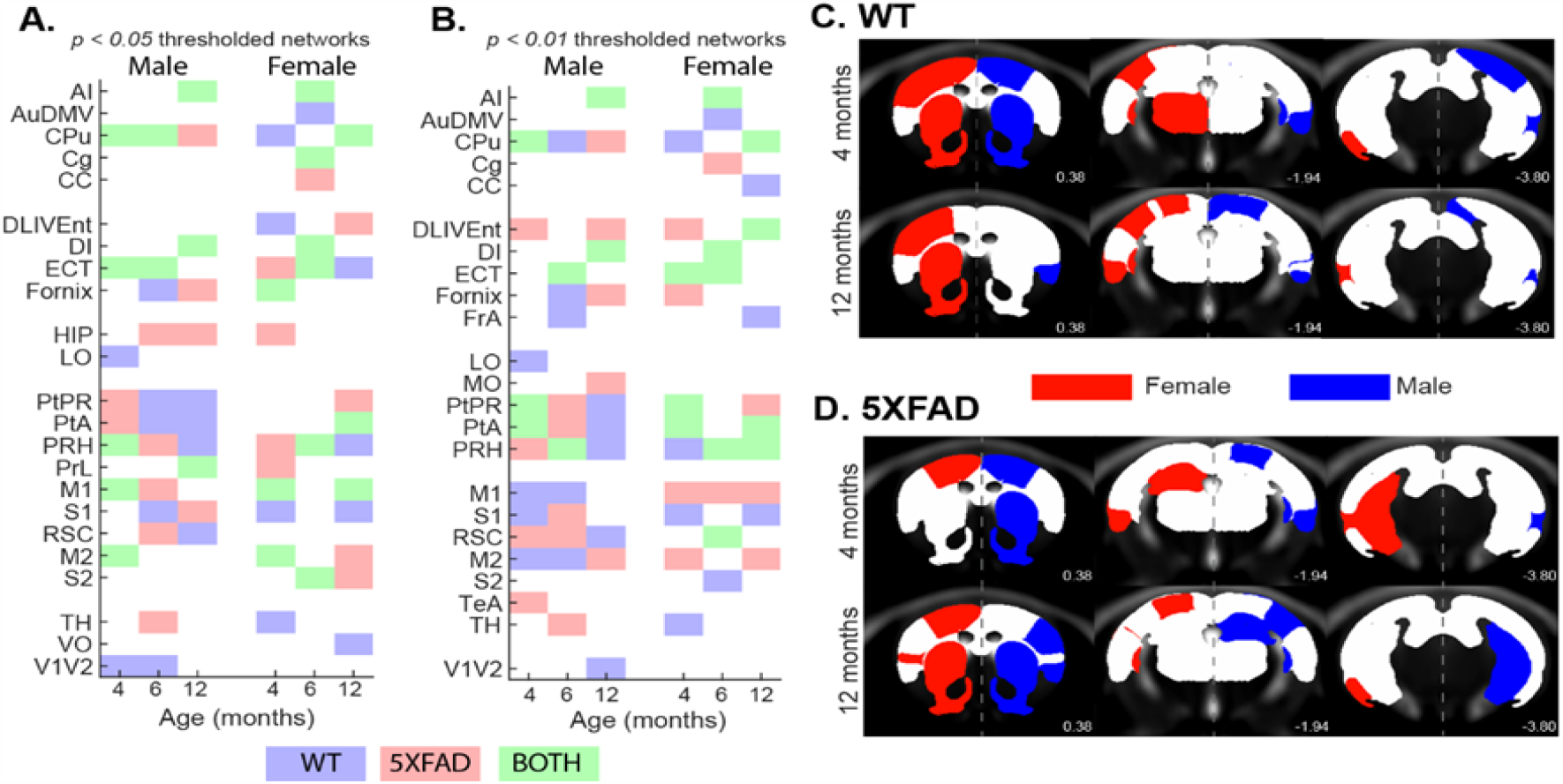
Covariance network degree hubs. Of the 27 nodes in the covariance networks, regions with the top 25% (7 regions) nodal degree were labeled as hubs for each group. Degree hubs are color coded based on whether they were identified in WT (blue), 5XFAD (red), or both (green) groups for *p* < 0.05 (A) and 0.01 thresholded networks. Spatial location of degree hubs from *p* < 0.05 networks for WT (C) and 5XFAD (D) groups are shown on a magnetic resonance image template for female (red) and male (blue) groups at youngest (top) and oldest (bottom) investigated age. Left/Right hemisphere separation by sex is for visualization purposes only. Images for 6-month age group degree hubs are shown in Figure S6. Region labels are in Table S1.

### Community/Modularity Structure of Metabolic Covariance

Leveraging a data-driven approach to derive a scale-independent consensus community structure from networks [31], we identified a consensus partition for each unthresholded group covariance network (Figure 5A). Partitions of WT networks for both sexes revealed a coarse consensus structure comprised largely of 2-3 communities. The exception was the 12-month female group, which had a finer scale consensus community structure of 7 communities. Relative to WT, the number of communities was greater in 5XFAD groups (with exception of 4-month male and 6-month female 5XFAD groups). Number of communities can be interpreted as an indicator of coherence among regions in the network, therefore, the observed higher community counts in 5XFAD groups demonstrate a disordered underlying metabolic covariance structure of those networks. Quantitative comparison of partition similarity among groups with adjusted mutual information (AMI) [33] showed diversity among consensus partitions, with no clear distinctions of age, sex, or genotype (Figure 5B). Co-classification (i.e., per node pair, the fractions of instances in which they were assigned to the same community) of consensus partitions within genotype revealed that WT groups had greater agreement for several regions, as seen in the Figure 5C co-classification matrix (left) and regional averages (right). Using the 4-month male WT partition as reference, mean SUVR values for the two modules were extracted and compared in a 3-way ANOVA (genotype, sex, and age as factors). For module 1 (Figure 5D), there was a main effect of age (F=9.9, *p* < 0.0005) as well as an age*sex interaction (F=4.68, *p* < 0.05). Post hoc tests revealed that 4- and 12-month groups were different from 6-month groups (Tukey-Kramer adjusted *p* < 0.05). No main effects or interactions were identified for module 2 (Figure 5E).

**Figure 5.**
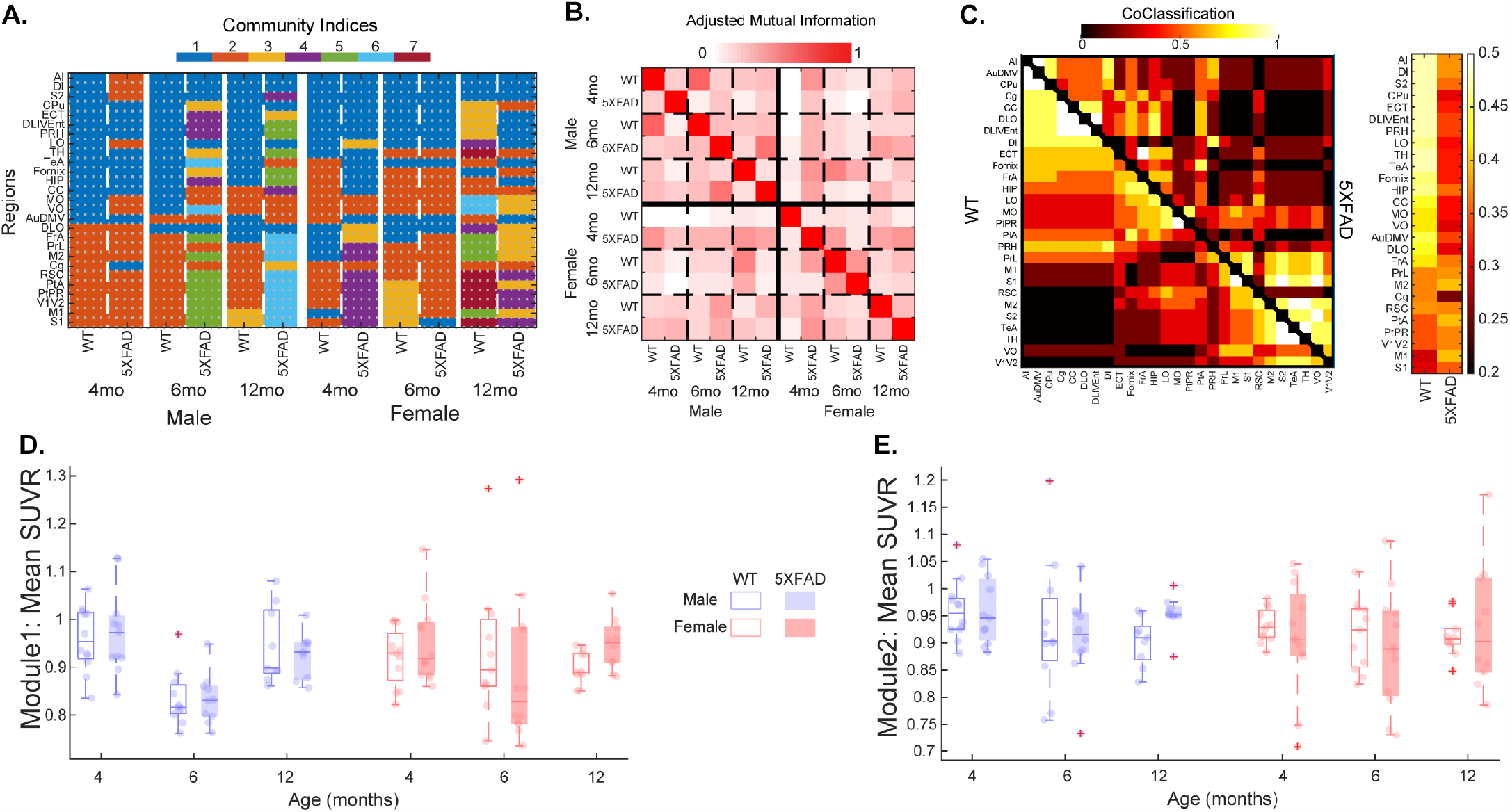
(A) Consensus modular structure for all groups (columns) with color indicating community assignment for each region (rows). (B) Adjusted mutual information matrix, quantifying the similarity of partitions among groups on a scale of zero (completely different) to one (identical). (C) Regional co-classification matrices computed from partitions of groups within genotype. WT data are in lower triangle and 5XFAD data are in the upper triangle. Average co-classification for each region within group is shown to the right of the matrix. (D-E) Using the 4-month male WT partition as reference, mean module SUVR values were extracted and are shown for (D) module 1 (corresponding to regions in blue in A for the 4-month male WT partition) and (E) module 2 (orange in the reference partition). Full names for row region labels in A can be found in Table S1.

## DISCUSSION

We used network connectivity analysis to understand group alterations in metabolic networks between and early-onset AD mouse model (5XFAD) and wild-type control groups (WT) at 4, 6, and 12 months in both sexes. Comparison of ^18^F-FDG SUVR covariance networks between WT and 5XFAD groups indicate altered interregional structure in the 5XFAD cohorts. While edge weight distributions were similar between the networks (Figure 1), multiple regions were found to differ in their covariance across genotype (Figure 2). Global and nodal network properties (i.e., density, degree, strength) showed a linear decrease with age in male cohorts and an inverted u-shape in female cohorts, with 5XFAD showing reductions in nodal degree and strengths, and differences in clustering of 5XFAD female group networks (Figure 3). These alterations in network structure suggest that AD pathology disrupts metabolic coupling, which is consistent with clinical evidence of glucose alterations in AD [34].

When pairwise relationships were assessed via statistical comparisons of individual edges between groups, multiple edges differed at an uncorrected *p* < 0.05, but did not survive FDR adjustment. Proper correction for multiple comparisons is an open topic in network neuroscience. Here, for a 27-region network, 351 unique tests are performed for each network pair. This makes it unlikely that individual edges would meet corrected significance. Correction methods such as Network-Based Statistics have been proposed as a more appropriate strategy [35]. In addition, edges contain shared information, so assumption of independence for parametric tests is not met. To properly assess group differences, we employed permutations-based testing (generating a null distribution of covariance networks by permuting group labels) to determine significance. Significant results from permutation testing as displayed in Figure 2 are information in implication a region or set of regions as altered in 5XFAD relative to WT, but we are cautious not to overemphasize individual connections, focusing on the whole rather than the parts.

Previous studies have reported both increases [23, 36] as well as decreases [8, 24] in ^18^F-FDG uptake of 5XFAD animals over the range from 7-12 months, with differences in younger mice. Given the lack of consensus, we hypothesized that interregional relationships deduced from covariance networks would better show AD-related alterations in glucose uptake measured with ^18^F-FDG PET. First, we found a significant difference in male vs. female weighted distributions in 4-month WT mice where the WT females showed more edges with stronger negative covariance, however, we did not observe this pattern in 4-month 5XFAD mice (Figure 1B). This lack of effect could be explained by the aggressive nature of the 5XFAD transgenic model. Indeed, Bouter et al. showed that early-age 5XFAD females exhibit lower brain metabolic uptake compared to controls. This global decrease likely overshadows any sex-dependent developmental differences in glucose metabolism [37]. Second, when comparing weighted distributions across genotype, we found a significant difference in edge weights (both positive and negative) between 12-month female WT and 5XFAD group networks (Figure 2F). These covariance networks are statistical constructs (i.e., correlations) that capture higher-order relationships beyond regional differences. Therefore, changes in the distributions of weights of these networks do not reflect hyper-or hypometabolism, but rather capture patterns shared by animals within a homogenous group. Altered ^18^F-FDG uptake can be reflective of reduced glucose utilization by the brain regions. These mechanisms are dependent on membrane transport proteins (i.e. GLUT1-5), which transport glucose across cellular membranes [34], and therefore it is plausible that regional expression of these proteins in the 5XFAD model contribute to observed alterations in metabolic covariance observed.

Global and nodal network properties from graph theory provide valuable insight into the organization and topology of networks. They can determine if networks are comparable, quantify the extent and magnitude of regional interrelationships, and assess characteristics of said networks [13, 38, 39]. The present characterization of metabolic covariance in the 5XFAD model of AD showed that in thresholded networks, network properties were sex and genotype dependent, as were the rates of change of said properties with respect to age. A recent study also reported age-related changes in metabolic covariance in young (2 months) and old (18 months) Sprague-Dawley rats, reporting edge-based global network differences, but not path-based ones [40]. Path-based measures utilized in Xue, Wu [40] were developed for structural connectivity networks, and their interpretations in functional connectivity or metabolic covariance networks are unclear. In their study, the shape of decline observed differed between groups, with networks of male groups showing a linear decline, while female groups showed and inverted u-shape. These patterns across age (Figure 3) were present in both WT and 5XFAD; however, male 5XFAD groups showed reductions at the earliest age of 4 months, while female 5XFAD groups showed increases at the latest time point (12 months). Increasing attention is being given in recent years to sex-specific differences in AD vulnerability and progression, noting faster decline greater pathology in women [41, 42], which is consistent with our findings.

Identifying nodes that show high connectivity/coupling to the rest of the network [39], known as hubs, can highlight key regions within networks. For metabolic covariance networks, hubs (here defined on degree and strength), are regions that show a similar highly connected ^18^F-FDG SUVR pattern across animals. There was a moderate degree of overlap among age, sex, and genotype. Notable hub nodes involved regions of the dorsal striatum, and entorhinal, parietal, and perirhinal cortices. Interestingly, the hippocampus reached our defined hub threshold only in the 5XFAD groups (Figure 4), while primary and secondary motor cortices were metabolic hubs only in the 4-month male WT cohort, 12-month male 5XFAD cohort, and as 4- and 12-month female cohorts of both genotypes. The strong covariance of motor cortical regions aligns with previous reports in mouse and rat models, which identified them as hubs in resting state functional MRI data [43, 44].

In addition to identifying potential hubs, we utilized data-driven modularity to identify community structure in the network of each group. Modules are characterized by greater within module connectivity than to between modules. Identifying the modular structure in a network is therefore an optimization problem; however, it can yield solutions of varying community size [45]. Here we employed multiresolution consensus clustering, which spans a space of module solutions across scale and statistically identifies a consensus modular partition [31]. For the twelve groups investigated here, modular partitions ranged between 2 and 7 modules. WT group networks had fewer modules, which qualitatively indicates a higher degree of similarity in regional metabolic uptake, while 5XFAD groups has a generally greater number of communities, indicating uptake stratification depending on functional grouping. Assessing overall similarity between all group pairs did not reveal any distinction of age, sex, or genotype. As a follow-up, nodal similarity in community assignment (i.e., co-classification) within genotype showed a pattern within WT groups that was disrupted in 5XFAD groups (Figure 5C). This difference further highlights the increased variability in metabolic covariance due to mutations in the 5XFAD model. Examining the community structure more closely (see Figure 5A), it is apparent that the covariance matrix of 12-month WT females is stratified based on function, with insular regions in module 1, auditory in module 2, rhinal in module 3, parietal in module 7, and orbitofrontal and limbic regions dispersed between the modules. The 12-month 5XFAD females, on the other hand, displayed similar all-in functional groupings in modules, but with more functional groups in each module. Module 1 contained all auditory, insular, and rhinal regions, module 2 had all basal ganglia and white matter, module 3 had the majority or orbitofrontal regions, and module 4 contained most of the parietal regions.

These findings extend our understanding of AD progression by modeling interregional relationships of glycolytic metabolism via ^18^F-FDG PET in 5XFAD mice; however, there are limitations to consider. First, this is a cross-sectional investigation with groups spanning age, sex, and genotype. Although the 5XFAD mice were from the same parental stock, and are therefore genetically identical, there is a possibility that rearing mice from different mothers, with unique microbiomes, could influence the ^18^F-FDG PET signals. Recent work has shown that C56BL6/J mice with identical rearing conditions have distinct gut microbiota, which has been reported to impact brain immune cell profiles and modulates cognitive and affective behavior in mice [46], and thus would be reflected in the uptake. To resolve these changes, one could construct within-subject metabolic networks from dynamic ^18^F-FDG PET studies. Second, the use of transgenic models, such as the 5XFAD, which over-express amyloid precursor protein, are known to show a strong sexual dimorphism resulting from the transgene expression being linked to the estrogen sensitive Thy1 promoter [47]. Previous and recent literature by Julliene et al. and Oblak et al. observed sexual dimorphism in the biology of 5XFAD mice [22, 48]. Julliene and colleagues found a pattern of blood-brain barrier leakage similar to the patterns of network characteristics observed in the current study (Figure 3). While males consistently showed deterioration in the blood-brain barrier over time, females deteriorated from 4-8 months but seemingly reversed deterioration from 8-12 months. This similar inverted paraboloid connects back to the idea that estrogen and the concentration of estrogen-related machinery (receptors, biomolecules, etc.) in the brain disproportionately affect structural integrity of the blood-brain barrier and metabolic uptake and thereby modify disease progression in the 5XFAD model of AD in a sex-dependent manner [49]. As the next generation of models are released from Model Organism Development & Evaluation for Late-Onset AD (MODEL-AD; www.model-ad.org), which do not employ transgenes, many of these issues will be overcome, thus clarifying the true role of sex in AD. The next generation of MODEL-AD will also utilize longitudinal studies as opposed to the parallel cohort methodology used in the present study. Longitudinal studies with multiple imaging time points will provide greater temporal resolution of metabolic uptake in mouse models of AD, the dynamics of which are integral to understanding AD pathophysiology. Another limitation arises from potential circadian rhythm variation in imaging. During the PET imaging phase, mice were brought into the well-lit imaging lab and PET images were acquired during their respective light cycle. However, imaging was conducted over the course of a 9-hour period and Krueger et al. [50] report that mice imaged in the later stages of their light cycle yield variation in metabolic uptake values. The current study did not consider these alterations, and future studies should be aware of potential circadian effects on metabolic uptake in ^18^F-FDG PET studies. Another limitation exists in the harmonization of the analysis methods for the primary data and lack of quantitative functional analysis of modularity. As described above, numerous analytical methods have been used in the literature to standardize uptake of ^18^F-FDG in both preclinical and clinical studies; however, each has significant limitations, and none can be used universally across species. In the case of SUVs, which make up the source information used for z-score transformed correlations, there is a possibility that the transformations themselves may artificially bias the results based on body weight differences alone. In the current study, we employed SUVR, using the cerebellum as a reference region, which is used in the AD Neuroimaging Initiative [51]; however, since the cerebellum is not a true “null” region, there is a chance that this normalization could introduce bias. Our results align with previous clinical findings [15], providing some level of assurance that the current approach is unbiased. In the current study, we gained functional insight from modular structures of MRCC-processed covariance matrices by using known functional relevance of brain regions and extrapolating the meaning of a module based on functional relevance. However, hierarchical relevance exists between regions, and a quantitative analysis that takes this into account, as well as the magnitude of metabolic uptake as recorded in regional SUVR, would be a valuable tool to implement to recognize the functional significance of modules in a more robust manner. Future studies could leverage the covariance framework to investigate the role of vasculature in conjunction with metabolism, as prior studies have found a neurovascular uncoupling effect that consistently follows a pattern in AD as a function of disease progression and aging [52, 53]. Finally, future studies could employ multi-modal analyses that generate and combine network data from PET (amyloid, tau, metabolism) and MRI (structural and functional) to obtain a more detailed understanding of the pathophysiology of AD.

## CONCLUSIONS

To our knowledge, this is the first comprehensive evaluation of metabolic covariance structure in the 5XFAD model of AD. The 5XFAD model mice show age related changes in interregional glucose uptake relationships that are altered relative to WT mice. Moreover, analysis of these data revealed that community structure of covariance networks was different across age and sex, with a disruption of interregional metabolic coupling in 5XFAD model of AD.

## DATA AVAILABILITY STATEMENT

The contributions presented in the study are publicly available. The imaging data are available via the AD Knowledge Portal: https://adknowledgeportal.org (permission was obtained for this material through a Creative Commons CC-BY license). The data, analyses, and tools are shared early in the research cycle without a publication embargo on secondary use. Data are available for general research use according to the following requirements for data access and data attribution (https://adknowledgeportal.org/DataAccess/Instructions).

## AUTHOR CONTRIBUTIONS

EJC, CB, RS, SCP, AAB, MV, and PRT contributed to conception and design of the study. SCP, AAB, RS and PRT performed studies and initial data analysis. EJC, RS, and PRT developed and tested the analysis software. EJC, CB, and PRT wrote the first draft of the manuscript. EJC, CB performed the statistical analysis. EJC, CB, and PRT wrote sections of the manuscript. All authors contributed to manuscript revision, read, and approved the submitted version.

## FUNDING

EJC - NIA T32 AG071444

PRT - NIA U54 AG054345

MV – NC for HPC CN00000013 CN1; ING PNC0000002_DARE; and WTDA 215747/Z/19/Z

## ACKNOWLEDGEMENTS

The authors would like to acknowledge the NIH/NIA and MODEL-AD consortium for making the data available through Open Science principles. The data for this study were funded through grants NIA T32AG071444 and U54AG054345. MV is supported by Italian National Center for HPC, BIG DATA AND QUANTUM COMPUTING (Project no. CN00000013 CN1), the Italian National Grant DIGITAL LIFELONG PREVENTION (Project no PNC0000002_DARE), and by Wellcome Trust Digital Award (no. 215747/Z/19/Z).

