## Supplemental Figures for "Brain Metabolic Network Covariance and Aging in a Mouse Model of Alzheimer’s Disease"

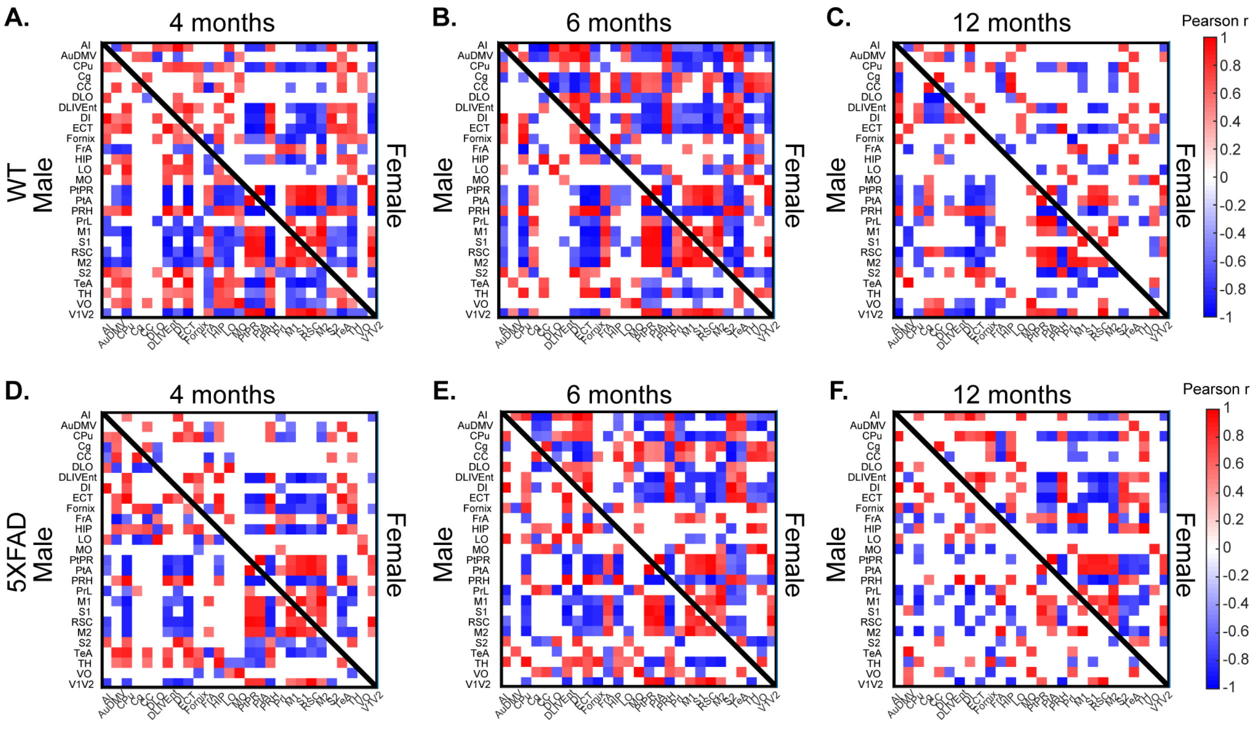


**Supplementary Figure 1.** Metabolic ($^{18}$F-FDG PET) covariance matrices for (A-C) wild-type (WT) and (D-F) 5XFAD groups. Matrices were thresholded at permutation $p < 0.05$ correlation significance. Matrices are Male (lower triangle of matrix) and Female (upper triangle of matrix) at 4 months (A, D), 6 months (B, E), and 12 months (C, F) old. Covariance was computed as Pearson correlation of *z*-scored regional standardized uptake value ratio values across animals within each group. Full names for row/column region labels can be found in Supplementary Table 1.

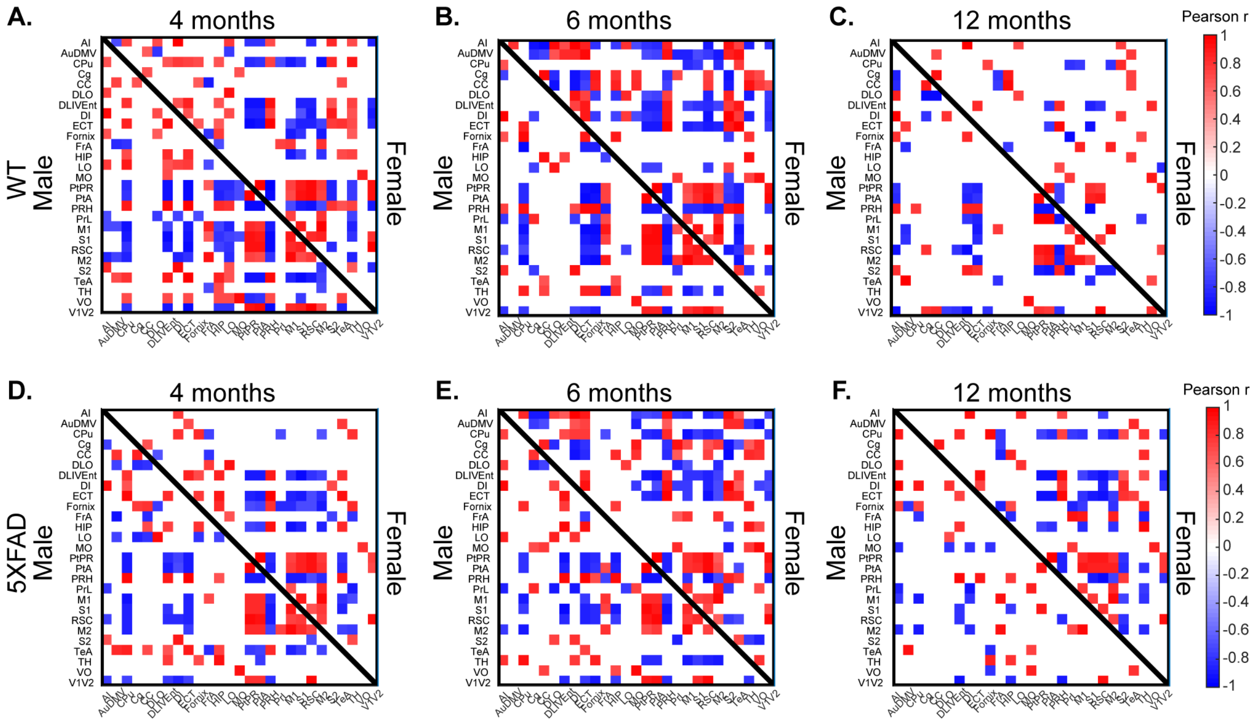


**Supplementary Figure 2.** Metabolic ([18]F-FDG PET) covariance matrices for (A-C) wild-type (WT) and (D-F) 5XFAD groups. Matrices were thresholded at permutation *p < 0.01* correlation significance. Matrices are Male (lower triangle of matrix) and Female (upper triangle of matrix) at 4 months (A, D), 6 months (B, E), and 12 months (C, F) old. Covariance was computed as Pearson correlation of *z*-scored regional standardized uptake value ratio values across animals within each group. Full names for row/column region labels can be found in Supplementary Table 1.

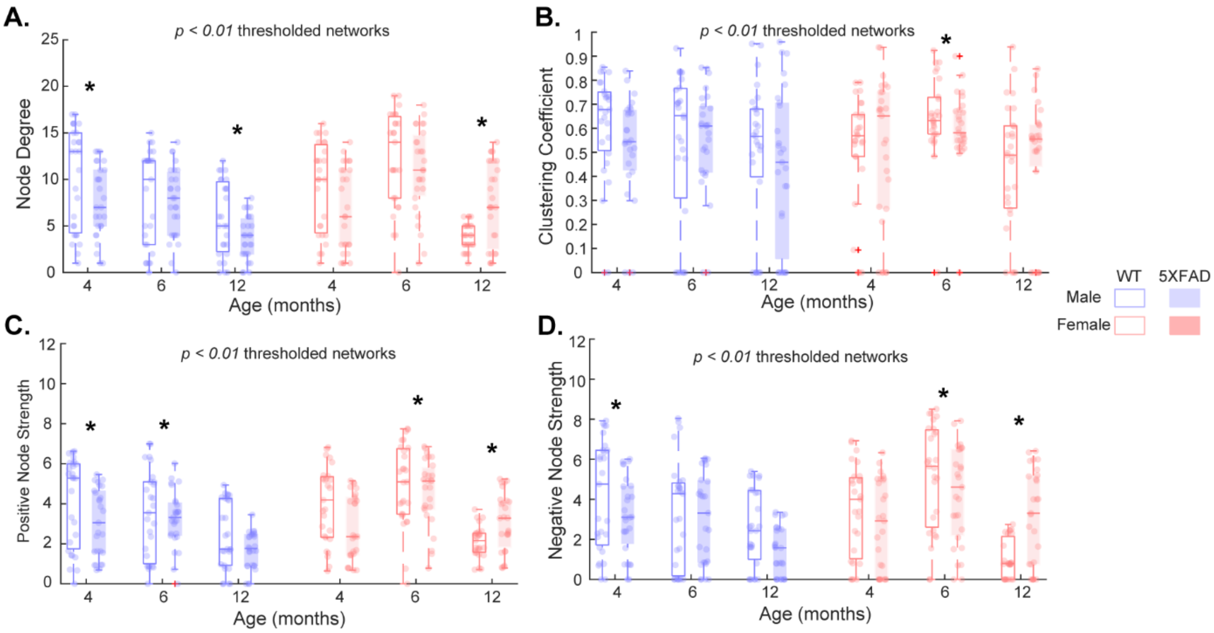


**Supplementary Figure 3.** Distributions of nodal properties of *$p < 0.01$* thresholded covariance networks for wild-type (WT) and 5XFAD groups at 4, 6, and 12 months old. *Denotes $p < 0.05$ 2-sample Kolmogorov-Smirnov test significance.

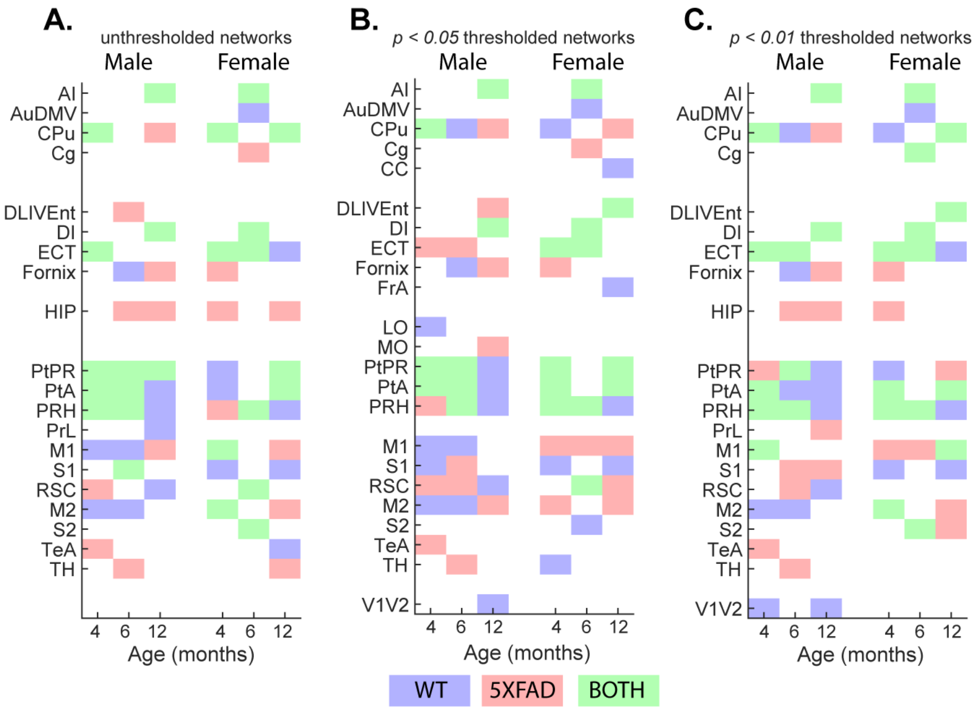


**Supplementary Figure 4**. Covariance network strength hubs. Of the 27 nodes in the covariance networks, regions with the top 25% (7 regions) nodal strength were labeled as hubs for each group. Strength hubs are color coded based on whether they were identified in WT (blue), 5XFAD (red), or both (green) groups for unthresholded (A) and $p < 0.05$ and 0.01 thresholded (B-C) networks. Spatial location of strength hubs for all ages as well as all thresholded network strength hubs are shown in Figure S5. Region labels are in Table S1.

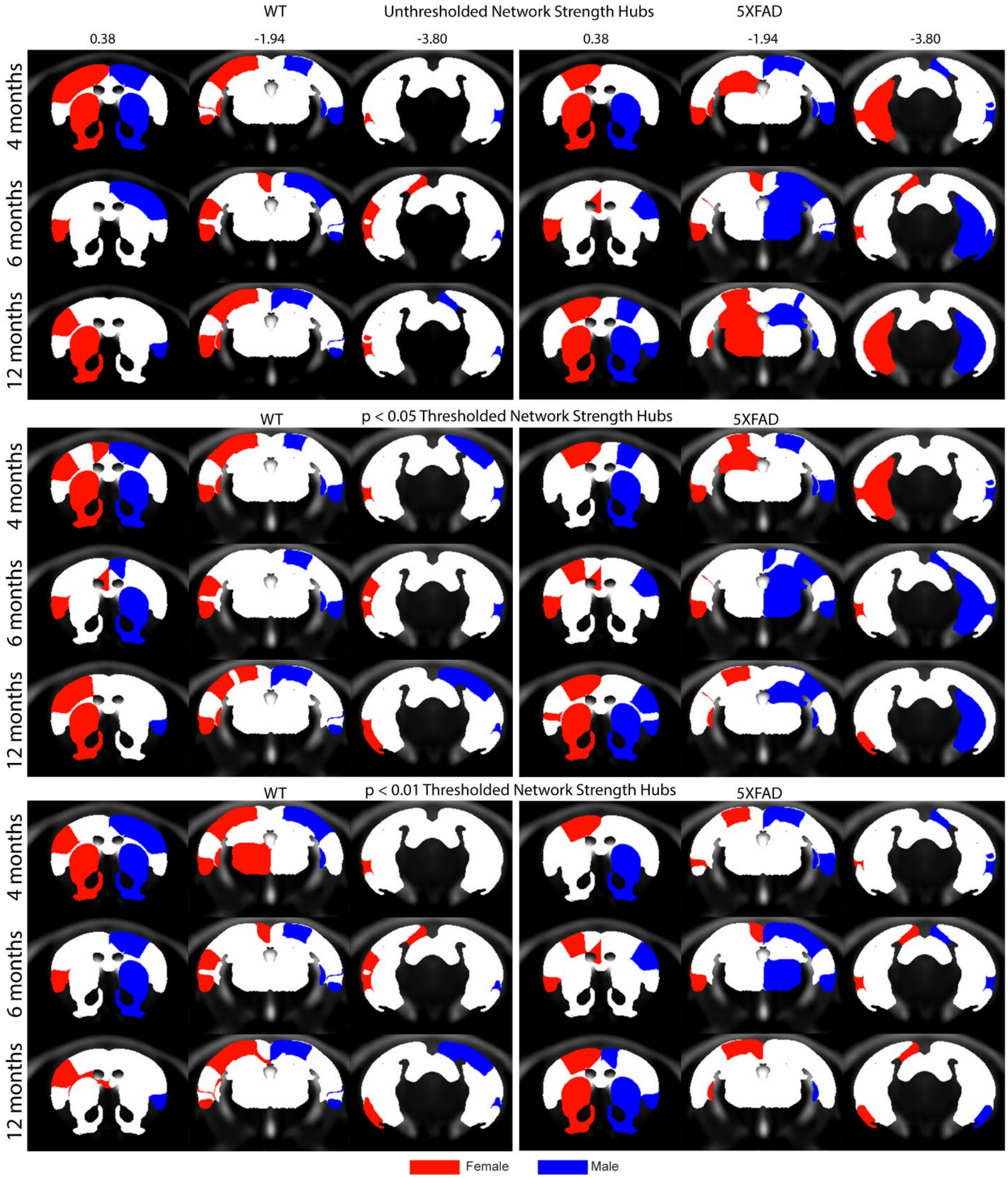


**Supplementary Figure 5.** Regions that were classified as hubs based on strength for all ages, groups, and thresholds.

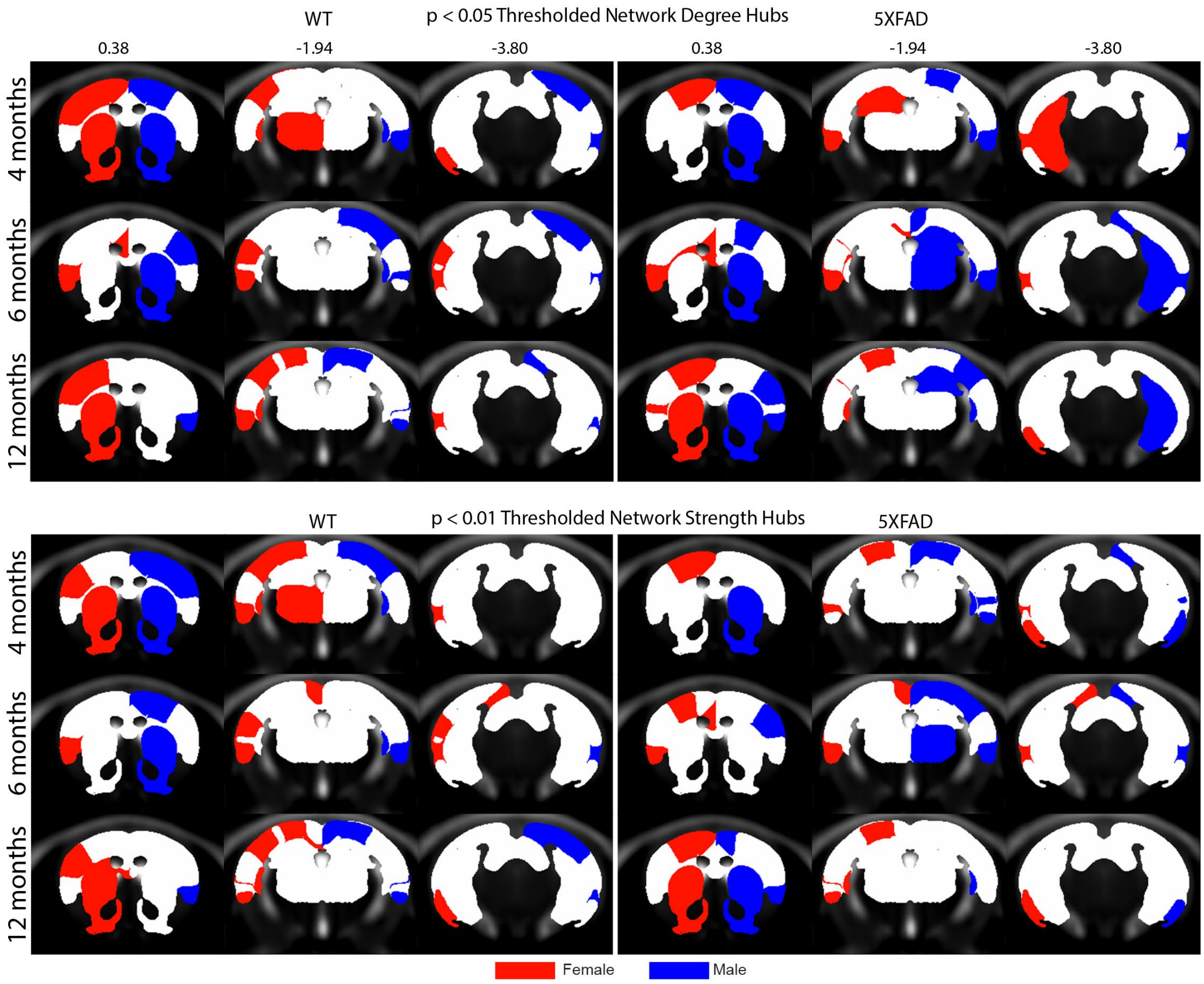


**Supplementary Figure 6.** Regions that were classified as hubs based on degree for all ages, groups, and investigated thresholds.

| Index | Region of Interest (ROI) Name | ROI Label |
| --- | --- | --- |
| 1 | Agranular Insular Cortex (Doral-Ventral) | AI |
| 2 | Auditory Cortex (Dorsal, Medial, Ventral) | AuDMV |
| 3 | Caudate Putamen (Dorsal Striatum) | CPu |
| 4 | Cingulate Cortex | Cg |
| 5 | Corpus Callosum | CC |
| 6 | Dorosolateral Orbital Cortex | DLO |
| 7 | Dorsintermed Entorhinal Cortex | DLIVEnt |
| 8 | Dysgranular Insular Cortex | DI |
| 9 | Entorhinal Cortex | ECT |
| 10 | Fornix | Fornix |
| 11 | Frontal Association Cortex | FrA |
| 12 | Hippocampus (CA1-CA3) | HIP |
| 13 | Lateral Orbital Cortex | LO |
| 14 | Medial Orbital Cortex | MO |
| 15 | Parietal Corext (Post-Rostral) | PtPR |
| 16 | Perietal Association Cortex (Lateral-Medial) | PtA |
| 17 | Perirhinal Cortex | PRH |
| 18 | Prelimbic Cortex | PrL |
| 19 | Primary Motor Cortex | M1 |
| 20 | Primary Somatosenory Cortex | S1 |
| 21 | Retrosplenial Dysgranular Cortex | RSC |
| 22 | Secondary Motor Cortex | M2 |
| 23 | Secondary Somatosenory Cortex | S2 |
| 24 | Temporal Association Cortex | TeA |
| 25 | Thalmus | TH |
| 26 | Ventral Orbital Cortex | VO |
| 27 | Visual Cortex (Primary and Secondary) | V1V2 |

**Supplementary Table 1.** Full names of network nodes (region of interest) as defined in the Paxinos and Franklin Atlas. Left and Right regions were averaged to yield a network of 27 regions.
